# DNA Repair Strategy Sets Transposon Mobilization Rates in *Caenorhabditis elegans*

**DOI:** 10.1101/2022.08.01.502254

**Authors:** Cindy Chang, Daniel J. Pagano, David D. Lowe, Scott Kennedy

## Abstract

Transposons are parasitic nucleic acids that threaten genome integrity in all cells. In the metazoan model organism *Caenorhabditis elegans*, DNA transposons are active in the soma where they are reported to exhibit mobilization rates ≅1000 fold higher than in germ cells. How and why DNA transposons might be so highly active in the *C. elegans* soma is a mystery. To better understand this question, we constructed reporter genes that label cells in which Tc1 has mobilized with fluorescent protein. The reporters recapitulate the known properties of DNA transposons in *C. elegans* and allow transposon activity to be monitored in intact, living animals. Using these reporters, we identify cytoplasmic and nuclear factors that limit transposition in the germline. Interestingly, none of these factors limit transposition in the soma. Rather, we identify a gene (*nhj-1*/*scb-1*), which we show is required for 99.9% of Tc1 mobilization events in somatic tissues, but does not influence mobilization in the germline. *nhj-1*/*scb-1* encodes a nematode-specific component of the non-homologous end joining (NHEJ) DNA repair machinery. Mutations in the other components of the NHEJ machinery (*cku-70*, *cku-80*, and *lig-4*) also suppress Tc1 mobilization in the *C. elegans* soma by ≅1000 fold. The data show that the use of NHEJ to repair transposon-induced DNA breaks in the soma dramatically increases the rate of transposon mobilization in this tissue. And because *C. elegans* germ cells use homology-based repair, and not NHEJ, to fix transposon-induced breaks, we propose that the 1000-fold difference in transposon mobility reported for the *C. elegans* soma and germline can, in large part, be explained by tissue-specific differences in DNA repair strategy.

**Author Summary:** Transposons are common parasitic genetic elements that threaten all genomes. For example, half of the human genome is made up of transposons. Transposon mobilization can disrupt gene function, causing disease, so transposon activity needs to be tightly regulated to prevent harm to the host. Transposons are typically less active in the soma than in the germline, because somatic transposition benefits neither host or transposon. Surprisingly, in the nematode model organism *Caenorhabditis elegans*, transposons are reported to be 1000-fold more active in the soma than the germline. Here, we develop a system to investigate transposon regulation in an intact live animal, and show that, in large part, tissue-specific differences in transposon activity in *C. elegans* is due to the use of different DNA repair pathways by these tissues, highlighting the importance of DNA repair strategy in determining outcomes of transposon excision events. Given that DNA repair factors have been linked to transposon regulation in other eukaryotes, we propose that DNA repair choice likely contributes to transposon mobilization in all eukaryotes.

## Introduction

Transposable elements are mobile genetic elements that threaten all eukaryotic genomes. Transposons are typically classified into two classes: The class I retrotransposons, which use an RNA intermediate and reverse transcription to replicate; and the class II DNA transposons, which mobilize via a direct DNA cut-and-paste or replicative transposition mechanisms [1]. The ability of class I and II transposons to mobilize and reproduce to high copy numbers presents challenges to all cells and organisms. For instance, newly mobilized transposable elements can alter gene expression programs, disrupt gene function, and increase the risk of chromosomal rearrangements. Therefore, a number of anti-transposon genome defense systems have evolved in all organisms, which limit the expression, replication, and mobilization of transposons. Such systems include small interfering small regulatory RNAs such as the small interfering (si)RNAs and PIWI-interacting (pi)RNAs, which silence transposon-encoded RNAs transcriptionally and post-transcriptionally in many animals [2, 3].

Transposable elements- and the remnants of ancient transposable elements-compose between 3-85% of all eukaryotic genomes [4]. About 12% of the genome in the metazoan model organism *C. elegans* is composed of transposons, with the most active and abundant of these being the class II Mariner DNA transposons [4, 5]. Most active Mariner elements in *C. elegans*, including Tc1-3 and Tc5, produce a single RNA that encodes a transposase enzyme that is responsible for the excision and re-insertion (cut-and-paste) of the element [5, 6]. Transposons that mobilize via cut-and-paste can expand within genomes via the following mechanisms. First, a transposon can mobilize from replicated DNA to non-replicated DNA during S phase, which can increase copy number [7]. Second, a DNA break caused by transposon excision can be repaired by homologous recombination (HR) using a homologous chromosome or sister chromatid as a template for repair. When this happens, HR repair restores the mobilized transposon back into its donor site and, if the original transposon inserts elsewhere in the genome, increases copy number [8]. This latter mechanism appears to operate in the *C. elegans* germline as animals made hemizygous for a Tc1 element exhibit a ≅100-fold increase in transposon removal from the donor site in germ cells, presumably because the homologous chromosome (in the hemizygous animals) lacks a transposon and, therefore, cannot template transposon restoration during repair [9]. In other words, the vast majority of transposon mobilization events occurring in germ cells are masked by HR-based repair, which re-copies an excised transposon back into its original position as a natural consequence of the repair process.

Forward genetic screens in *C. elegans* have identified genes and pathways that limit transposon mobilization in the *C. elegans* germline [10–12]. Hereafter, we use the terms “mobilized or mobilization” to refer to transposons no longer present at their original chromosomal position. *rde-3*/*mut-2* (henceforth, *rde-3*) encodes a nucleotidyltransferase that silences transposons by appending long stretches of alternative U and G nucleotides to the 3’ termini of Tc1 RNAs [13, 14]. 3’ UG repeats (or poly (p)UG tails) recruit an RNA dependent RNA Polymerase (RdRP) enzyme that uses the pUGylated RNA as a template to synthesize antisense siRNAs, which bind Argonaute proteins to inhibit Tc1 transposase expression and, therefore, mobility [13]. Another gene required for transposon silencing in *C. elegans* is *mut-16* which encodes a low-complexity protein that helps recruit transposon-regulating factors, such as RDE-3, to a sub-compartment within germ granules to help silence Tc1 RNA [12,15,16]. Interestingly, RDE-3 does not regulate Tc1 activity in the soma, suggesting that the germline and the soma may possess different systems for controlling their genomic parasites [10].

In many organisms in which there is clear distinction between germline and soma, transposable elements are more active in the germline than in the soma. For instance, in *Drosophila*, *P* elements transpose in the germline, but are largely inactive in the soma because functional transposase protein is only produced in the germline [17–19]. Similarly, the non-long terminal repeat (non-LTR) LINE-1-like retrotransposon *I* factor is active in the germline, but not in the soma of flies [20]. And in mice, non-LTR retrotransposon LINE1 elements are active in germ cells and early stage embryos, but are much less active in somatic tissues, with the possible exception of neuronal progenitor cells [21–23]. It has been suggested that the reason transposons are typically inactive in the soma is because neither the host nor the transposon will benefit from transposition in the soma [24]. More specifically, somatic transposition causes genetic damage and, therefore, disadvantages the host, as evidenced by the fact that transposon activity in people leads to disease [25]. Additionally, somatic transposition does not benefit the transposon, because any increase in transposon number accompanying transposition is not inherited. Therefore, evolution has likely selected for transposons (and hosts) that show low levels of transposition within somatic tissues when compared to the germline, where transposition will at least benefit the transposon [24]. A striking deviation from this paradigm comes from *C. elegans* where Mariner DNA transposons are reported to be ≅1000-fold more active in the soma than in the germline [26, 27]. How or why Tc1 might be so highly active in the *C. elegans* soma is not known.

Here we report the development of reporter genes for visualizing Tc1 mobilization in the soma and germline of living *C. elegans*. We use the reporters to identify novel cytoplasmic and nuclear factors that limit transposon mobilization in the germline, but not in the soma. We also identify a gene *nhj-1/scb-1* that regulates transposon mobilization in the soma, but not in the germline. We use this mutant to establish that the use of NHEJ to repair transposon-induced chromosome breaks in the soma results in a 1000-fold increase in transposon mobilization rates in this tissue. And because *C. elegans* uses HR to repair transposon-induced DNA breaks in the germline, and HR (but not NHEJ) masks transposon excision events, we propose that the radically different levels of transposon activity reported for the *C. elegans* soma and germline are not due to differing levels of transposon activity *per se* but, rather, to the use of different DNA repair pathways to fix transposon-induced breaks in these tissues.

## Results

### Design of reporter genes to visualize transposon mobilization in the *C. elegans* soma and germline

*C. elegans* Tc1 is a Mariner class DNA transposon which is mobile in the *C. elegans* soma and germline [6]. To explore the biology of transposons in an intact animal system, we engineered a reporter gene under the control of the ubiquitous promoter (*eft-3*p) to drive nuclear expression of mScarlet in all cells of the *C. elegans* soma and germline. A Tc1 element was then inserted into the promoter of *eft-3*p*::mScarlet* (Fig 1A) with the expectation; 1) that the transposon would interfere with mScarlet expression; 2) that transposon mobilization would permit mScarlet re-expression in just those cells in which Tc1 had mobilized; and finally 3) because *C. elegans* is transparent, these reporters would allow us to monitor Tc1 activity in all cells and during all stages of development of an intact animal model system. Note that Tc1 mobilization in the germline would be expected to result in animals that express fluorescent protein in all cells of the germline and soma and this expression state should be heritable. Somatic mobilization, on the other hand, would be expected to result in animals expressing fluorescent protein in one or more somatic cells and the expression pattern would not be inherited. We used CRISPR/Cas9 to insert the *Tc1::mScarlet* reporter into the genome. A control mScarlet reporter gene lacking the Tc1 element was also inserted into the identical chromosomal site. Fluorescent imaging revealed that, as expected; 1) mScarlet was expressed in all cells in *eft-3*p*::mScarlet* animals; and 2) the addition of a Tc1 element inhibited mScarlet expression in all cells (Fig 1B).

**Fig 1.**
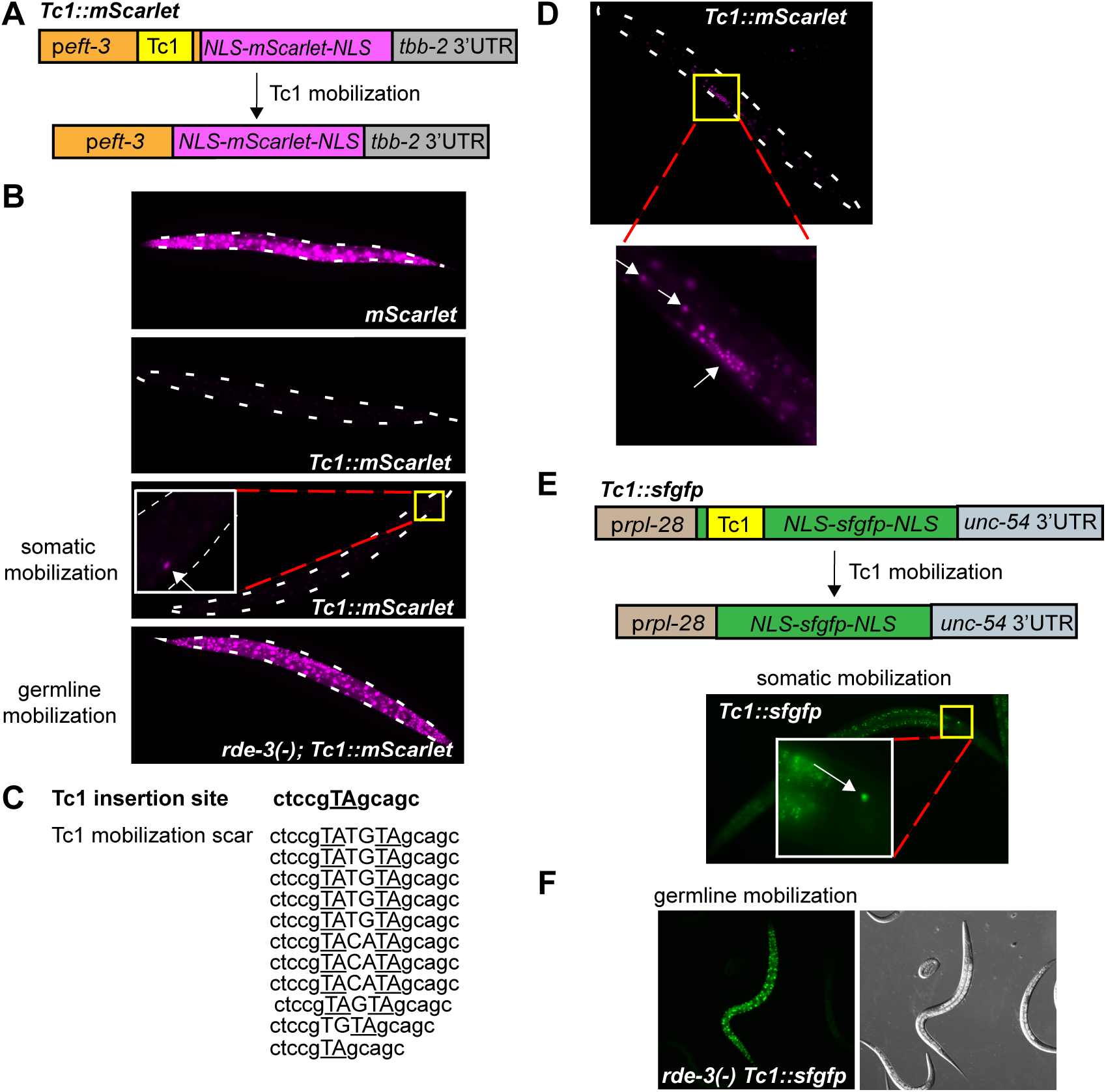
Transposon reporter genes detect Tc1 mobilization events in intact animals. (A) Schematic of the *Tc1::mScarlet* reporter gene engineered to monitor Tc1 mobilization. Tc1 was inserted within the promoter, with the expectation that Tc1 mobilization from the promoter would allow mScarlet expression. Nuclear localization signal (NLS). (B) Fluorescence micrographs of *eft-3*p*::mScarlet* (*mScarlet*), and *eft-3*p*::Tc1::mScarlet* (*Tc1::mScarlet*) animals. Somatic mobilization; an animal with one somatic cell expressing mScarlet is shown. A magnification of this cell is also shown and is indicated by an arrow. Germline mobilization; a germline mobilization event in an *rde-3(-)* animal is shown. mScarlet expression is observed in all cells of the germline and soma. Throughout the course of this work, no germline mobilization events were ever observed in wild-type *Tc1::mScarlet* animals. (C) Sanger sequencing of Tc1 mobilization events from DNA derived from *Tc1::mScarlet* animals possessing one or more fluorescent somatic cells (see Materials and Methods for details). (D) Fluorescence micrographs of Tc1 mobilization in the M-lineage of a *Tc1::mScarlet* animal. Arrows indicate cells with bright mScarlet fluorescence, indicating Tc1 mobilization. (E) Schematic of *Tc1::sfgfp* reporter and a fluorescence micrograph of a somatic Tc1 mobilization event in a *Tc1::sfgfp* animal (bottom panel). *Tc1::sfgfp* was designed such that a +4 transposition scar, which is typical for Tc1 mobilization events, would restore the correct reading frame to *sfgfp.* Nuclear localization signal (NLS). Magnification of the yellow boxed area is shown, with a sfGFP expressing cell indicated by an arrow. (F) An *rde-3(-)* animal expressing sfGFP in all cells of the soma and germline, indicating germline mobilization. Throughout the course of this work, no germline mobilization events from wild-type *Tc1::sfgfp* animals were ever observed in wild-type animals.

### Tc1 reporter genes recapitulate known transposon behaviors

We next asked if the *Tc1::mScarlet* reporter gene recapitulated the known properties of Tc1 and, therefore, could be used to study transposon regulation in a living animal. [Note, the *Tc1::mScarlet* reporter gene will not report on the insertion (paste) step of transposition.] Endogenous Tc1 transposons exhibit a very low rate of mobilization in wild-type *C. elegans* germ cells (<5×10^-8^) [28, 29]. We found that Tc1 mobilized from *Tc1::mScarlet* in the germline at a similarly low rate. *Tc1::mScarlet* germline mobilization rates were determined by growing animals to high density over several generations and then determining the percentage of growth plates containing one or more animals expressing mScarlet in all cells of the soma and germline. A Poisson distribution method (see Materials and Methods) was then used to set a maximum mobilization rate [10, 28]. This approach set a germline mobilization rate for *Tc1::mScarlet* animals at <2.8×10^-7^, which is similar to published mobilization rates for Tc1 elements inserted into endogenous *C. elegans* genes (<5×10^-8^) (Table 1) [28, 29]. Five additional lines of evidence indicate that the *Tc1::mScarlet* reporter gene faithfully recapitulates Tc1 biology in the *C. elegans* soma and germline.

**Table 1.** Germline Tc1 mobilization rates determined using the Tc1::mScarlet reporter. Animals heritably expressing mScarlet in all cells of the germline and soma were scored as positive for germline mobilization. For some genetic backgrounds (rde-3(ne3370), mut-16(pk710), and deps-1(bn124), in which mobilization rates were high, the rate of germline mobilization rate was determined by scoring plates with F1 broods. For genetic backgrounds with lower rates of germline mobilization, plates were scored with F2/3 broods (znfx-1(ne4583) and nrde-2(gg873)). For genetic backgrounds where the rate of germline mobilization was so low that no mobilization events were ever observed (demarcated with *), the Poisson distribution method was used to set a maximum rate of germline mobilization based on scoring of a large number of F2/3 animals (see Materials and Methods). Data are aggregates from scoring from at least 3 independent experiments, per genotype. ND, not determined.

| Genotype | # plates scored | # animals/ plate (F1) | # animals/ plate (F2/F3) | # plates with animal(s) expressing mScarlet in all cells | Germline mobilization rate |
| --- | --- | --- | --- | --- | --- |
| *Tc1::mScarlet* * | 122 | ND | 29120 | 0/122 | $<2.8 \times 10^{-7}$ |
| *rde-3(ne3370); Tc1::mScarlet* | 81 | 322 | ND | 54/81 | $3.4 \times 10^{-3}$ |
| *mut-16(pk710); Tc1::mScarlet* | 32 | 416 | ND | 19/32 | $2.2 \times 10^{-3}$ |
| *mut-16(mg461); Tc1::mScarlet* * | 34 | ND | 29958 | 0/34 | $<1 \times 10^{-6}$ |
| *deps-1(bn124); Tc1::mScarlet* | 33 | 1183 | ND | 14/33 | $4.7 \times 10^{-4}$ |
| *znfx-1(ne4583); Tc1::mScarlet* | 36 | ND | 22513 | 10/36 | $1.4 \times 10^{-5}$ |
| *nrde-2(gg873); Tc1::mScarlet* | 30 | ND | 22026 | 5/30 | $8.3 \times 10^{-6}$ |
| *nhj-1(gg875); Tc1::mScarlet* * | 39 | ND | 26309 | 0/39 | $<9.9 \times 10^{-7}$ |

First, Tc1 is reported to be ≅1000 fold more active in the *C. elegans* soma than in the germline [10,26,27]. We observed that 23% of *Tc1::mScarlet* animals expressed high levels of mScarlet in one or more somatic cells, suggesting that Tc1 had mobilized in these cells (Fig 1B). The somatic rate of *Tc1* mobilization was determined by dividing the total number of animals possessing one or more mScarlet positive cells (329) by the total number of somatic cells scored (≅1.33×10^6^) (Table 2). This analysis sets the somatic mobilization rate at ≥2.5 x10^-4^. Thus, the rate of Tc1 mobilization from *Tc1::mScarlet* is ∼900-fold higher in the soma than in the germline (≥2.5 x10^-4^ vs. <2.8×10^-7^), which is consistent with previous reports [26].

**Table 2.** Somatic Tc1 mobilization events subcategorized by the number of somatic cells expressing mScarlet. Somatic Tc1 mobilization events were detected by mScarlet fluorescence in somatic cells (indicated as (+)). The rate of somatic mobilization was determined as follows: Number of animals with (+) cells (329)/((total number of animals scored (1333)) x (# somatic cells/animal (≅1000)). Note this number may be an underestimate as it is possible that we did not score all somatic mobilization events. ^a^15 out of these 233 animals had (+) signal in two neighboring gut nuclei. These were categorized as 1 (+) cell animals as some/many of these animals likely underwent a single transposition event with mScarlet localizing to both nuclei of the gut cells, which are binucleated and share a cytoplasm.

| # Animals scored | # Animals with 1 (+) cell | # Animals with 2 (+) cells | # Animals with >2 (+) cells | # Animals with M lineage (+) cells | Somatic Tc1 mobilization rate |
| --- | --- | --- | --- | --- | --- |
| 1333 | 233<sup>a</sup> | 77 | 16 | 3 | $\geq 2.5 \times 10^{-4}$ |

Second, 78% of the time after excision, endogenous Tc1 elements, which always insert into TA dinucleotides, leave behind a four base pair TATGTA or TACATA mobilization footprint [27]. We cloned and sequenced eleven independent *Tc1::mScarlet* mobilization events and in 8/11 cases (73%), we observed a TATGTA or TACATA four base pair insertion at the TA dinucleotide formerly occupied by Tc1 (Fig 1C), which is consistent with previous reports [27].

Third, Tc1 is known to occasionally mobilize from the *unc-54* gene in all/most cells of the M-lineage during embryonic development [26, 30]. We observed *Tc1::mScarlet* animals that expressed mScarlet in all cells of the M-lineage (0.22% of animals observed, Fig 1D and Table 2), which is consistent with previous reports and, also, confirms that Tc1 can mobilize in mitotically dividing cells. It is worth noting, however, the majority of somatic mobilization events that we observed in *Tc1::mScarlet* animals (71%), occurred in animals possessing only one detectable mScarlet positive cell (Table 2), suggesting that most Tc1 mobilization events in the *C. elegans* soma occur in post-mitotic cells.

Fourth, we constructed a second, independent Tc1 reporter gene in which a Tc1 element was inserted into the coding sequence of a superfolder green fluorescent protein (*sfgfp)* gene driven under the control of a distinct, ubiquitous promoter (*rpl-28*p) (Fig 1E). The reading frame of *Tc1::sfgfp* was engineered such that mobilization of Tc1 from *Tc1::sfgfp* would permit sfGFP translation (Fig 1E). *Tc1::sfgfp* behaved like *Tc1::mScarlet*, exhibiting biological properties previously ascribed to endogenous Tc1 elements (Figs 1E and 1F).

Fifth, genetic analyses to be described below will show that the Tc1 elements within *Tc1::sfgfp* and *Tc1::mScarlet* are regulated by factors previously shown to regulate endogenous Tc1 elements in the germline (see below).

Thus, the Tc1 reporter genes can be used to explore transposon biology in an intact animal system.

### Identification of factors silencing Tc1 in the germline

Genetic screens in *C. elegans* have identified genes, such as *rde-3* and *mut-16*, which are required for limiting Tc1 mobility in the germline. [Note that data presented below will show that mobilization rates can be influenced by excision frequency as well as DNA repair pathway choice.] *rde-3* encodes a poly(UG) nucleotidyltransferase [10,13,14] and *mut-16* encodes a low complexity poly Q/N protein MUT-16 [11]. Both RDE-3 and MUT-16 localize within germ cells to non-membrane enclosed cytoplasmic organelles termed the *Mutator* foci, where they contribute to the biogenesis of endogenous (endo) small interfering (si)RNAs, which are thought to negatively regulate the Tc1 transposase mRNA, thereby preventing Tc1 transposition [15,31– 33]. We found that *rde-3(-);Tc1::mScarlet* animals exhibited a >1000 fold increase in Tc1 mobilization from *Tc1::mScarlet* in the germline (Fig 2A and Table 1). Similarly, *mut-16(pk710)* animals, which lack MUT-16 [11, 33], exhibited a >1000-fold increase in the rate of Tc1 mobilization in the germline (Fig 2A and Table 1). The data confirm previous reports showing that RDE-3 and MUT-16 silence DNA transposons in the germline and, importantly, establish that *Tc1::mScarlet* is regulated by pathways similar to those regulating endogenous Tc1 elements.

**Fig 2.**
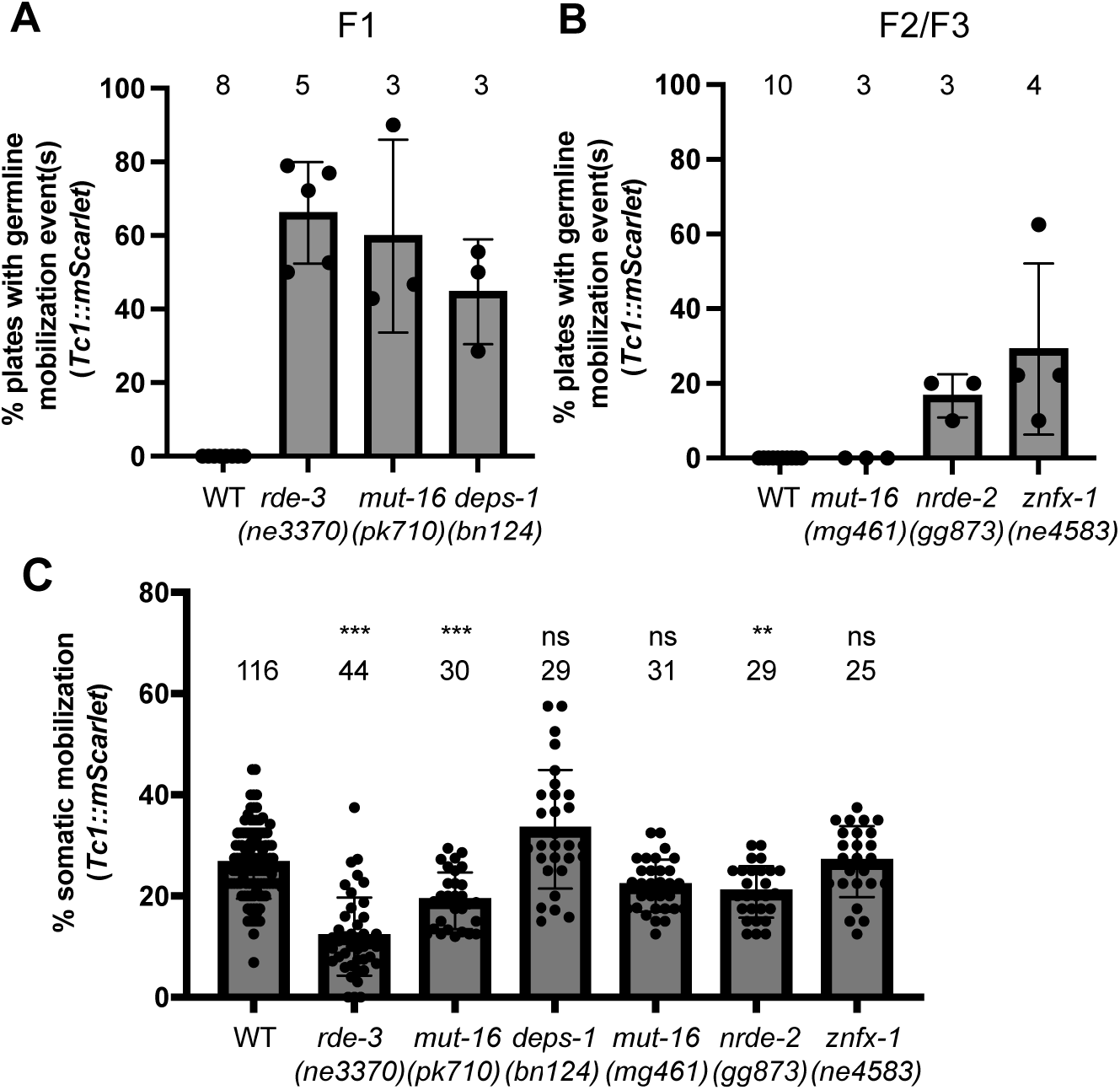
Germline anti-transposon defense systems do not silence transposons in the soma. (A-B) Animals of the indicated genotypes expressing *Tc1::mScarlet* were singled onto individual plates and the (A) F1 progeny and (B) F2/F3 progeny were scored for germline mobilization events (mScarlet expression in all cells). (A-B) Percentage of plates showing one or more germline mobilization events in the indicated generation is shown. (C) Progeny of a single *Tc1::mScarlet* expressing animal of the indicated genotype were scored, blinded to genotype, to determine the percentage of animals expressing mScarlet in one or more somatic cells. Data points represent independent lineages scored and are from at least three independent experiments scored on three different days. (A-C) Error bars are standard deviations of the mean. Alleles tested include (A)(C) *rde-3(ne3370)*, *mut-16(pk710)*, *deps-1(bn124)*; (B)(C) *mut-16(mg461)*, *nrde-2(gg873)*, *znfx-1(ne4583)*. ns, not significant, **p<0.01; ***p<0.001 (Kruskal-Wallis test with Dunn’s test for multiple comparisons).

We used *Tc1::mScarlet* to ask if other siRNA-related genes and pathways might coordinate with *Mutator* foci to silence transposons in the germline. *Mutator* foci localize near the outer membrane of germ cell nuclei juxtaposed to nuclear pores and directly adjacent to two other perinuclear foci termed the P granule and the Z granule [15,16,34–36]. Animals lacking DEPS-1, which is a low-complexity protein that localizes to P granules and is needed for P granule formation [16, 37], exhibited a large increase in germline Tc1 mobilization (Fig 2A and Table 1). And animals lacking the helicase ZNFX-1, which localizes to Z granules where it binds mRNAs undergoing siRNA-based gene silencing [16], exhibited a moderate increase in germline Tc1 mobilization, which became evident after lineages were grown to high densities for two to three generations (Fig 2B and Table 1). The data suggest that the perinuclear germ cell foci coordinately prevent transposon mobilization, perhaps by concentrating siRNA pathway proteins near nuclear pores to surveille mRNAs exiting nuclei for signs of aberrancy [35,36,38,39]. In addition to these cytoplasmic functions, *C. elegans* siRNAs also regulate gene expression within nuclei by modifying chromatin states and inhibiting transcription elongation (termed nuclear RNAi) [40, 41]. We asked if the nuclear RNAi pathway might contribute to transposon silencing. Indeed, animals lacking NRDE-2, which is an RNA binding protein required for nuclear RNAi [42] exhibited a moderate increase in germline Tc1 mobilization, which was evident after two to three generations of growth (Fig 2B and Table 1). We conclude that the cytoplasmic and nuclear branches of the endogenous RNAi system limit transposon mobilization in the *C. elegans* germline.

### Germline anti-transposon systems do not regulate Tc1 in the soma

NRDE-2, MUT-16, and RDE-3, which silence transposons in the germline (see above), are expressed and/or are active in both the germline and the soma [13,33,42]. Therefore, we asked if any of the inhibitors of germline mobilization described above might also inhibit mobilization in the soma. Surprisingly, presumed null mutations in *rde-3*, *deps-1*, *znfx-1*, or *nrde-2* did not increase Tc1 mobilization in the soma (Fig 2C) as they did in the germline (Figs 2A and 2B). Note that *rde-3, mut-16* and *nrde-2* mutants actually exhibited a subtle (< 2x) yet statistically significant reduction in Tc1 mobility in the soma (Fig 2C). Reasons why these factors might normally promote, albeit subtly, transposition in the soma are not known. The *mut-16(pk710)* allele ablates MUT-16 function in both the soma and germline while the *mut-16(mg461)* allele deletes a portion of the *mut-16* promoter, which abolishes MUT-16 function in the soma, but not the germline [33]. While *mut-16(pk710)* animals showed elevated rates of Tc1 mobilization in the germline (Fig 2A), somatic mobilization rates were unaffected (Fig 2C). The soma specific allele *mut-16(mg461)* failed to alter mobilization rates in either the germline or the soma (Figs 2B and 2C). Together, the data show that the known germline anti-transposon systems do not silence Tc1 mobilization in the soma, despite some of the components of these pathways being present and active in this tissue.

### Identification of a polymorphism that suppresses Tc1 mobilization in the soma

We next set out to explore how transposons might be regulated in the soma. We were surprised to find that introduction of *Tc1::mScarlet* into the identical chromosomal position of two ostensibly isogenic wild-type strains of *C. elegans* resulted in animals exhibiting radically different rates of Tc1 mobilization (Fig 3A). Hereafter, we refer to ostensibly wild-type *Tc1::mScarlet* strains with low soma Tc1 mobilization rates as *Tc1::mScarlet* (Low, L) and ostensibly wild-type *Tc1::mScarlet* strains with high soma Tc1 mobilization rates as *Tc1::mScarlet* (High, H). Genetic crosses between H and L animals resulted in progeny exhibiting H and L phenotypes in a pattern suggestive of a recessive mutation being present in L animals and causing the L phenotype (S1 Fig). The data suggests that the rate at which Tc1 mobilizes in somatic cells is under genetic control and that a background polymorphism present in some ostensibly wild-type strains of *C. elegans* dramatically alters the rate at which Tc1 will mobilize in the soma.

**Fig 3.**
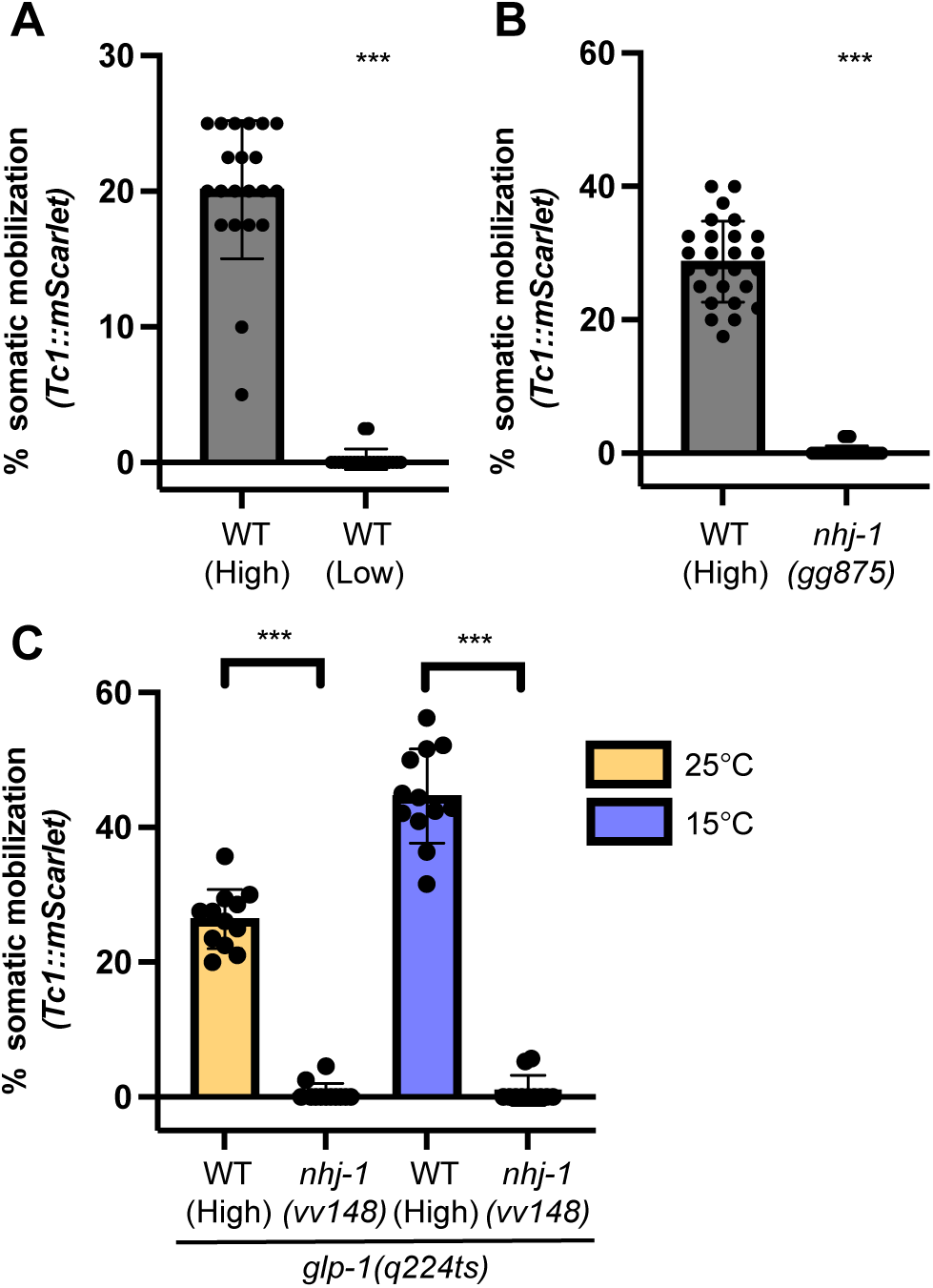
*nhj-1* mutations suppress Tc1 mobilization in the soma. (A) Two ostensibly wild-type (WT) strains show dramatically different levels (High or Low) of somatic Tc1 mobilization. (B) A putative *nhj-1* null allele (*gg875*) suppresses somatic Tc1 mobilization in wild-type (High) animals. (C) NHJ-1 regulates Tc1 mobilization in the soma. (A-C) Y axis, % of *Tc1::mScarlet* animals possessing one or two somatic cells expressing mScarlet indicative of Tc1 mobilization from *Tc1::mScarlet*. (A-B) Progeny of a single *Tc1::mScarlet* animals of the indicated genotypes were scored, blinded to genotype, for the percentage of animals in each lineage expressing mScarlet in one or more somatic cells. Each data point represents a lineage. Data-points are from at least three independent experiments scored on three different days. Error bars are standard deviations of the mean. ***p<0.001 (Mann-Whitney test). (C) *glp-1(q224ts)* animals lack most germ cells when grown at 25°C. Animals were bleach-synchronized and raised at 15°C or 25°C prior to scoring. A population of animals of the indicated genotypes were scored at larval stage 4 after synchronization, blinded to genotype, for the percentage of animals in each population expressing mScarlet in one or more somatic cells. Each data point represents an independent population. Data-points are from at least three independent experiments scored on three different days. Error bars are standard deviations of the mean. ***p<0.001 (Mann-Whitney test).

### NHJ-1 regulates Tc1 mobility in the soma

We sought to identify the mutation responsible for regulating Tc1 in the soma. Data presented below will show that the causal mutation is in the gene *nhj-1*/*scb-1* and that the mutant allele is present in L animals. For clarity we will henceforth refer to the gene as *nhj-1.* [Note, images and data presented in Figs 1-2 and Tables 1-2 were generated with H animals, which lack the *nhj-1* mutation, unless otherwise indicated.] To positionally map *nhj-1*, we crossed wild-type (L) animals to a strain from the Million Mutation Project, which exhibited an H phenotype [43]. We isolated 75 recombinant F2 progeny exhibiting an L phenotype and used sequence polymorphisms in the two strains to map *nhj-1* to Chr V, between 9.4 and 12.6 Mb (S2 Fig). We conducted whole genome sequencing of L and H animals. Variant calling programs (samtools/bcftools in the Mutation Identification in Model Organism Genomes (MiModD) software package [44]) failed to identify DNA alterations between H and L animals within the mapping interval. However, manual inspection of the data identified a small region within the *nhj-1* gene, located at 11.1 Mb on Chr V, in which sequencing coverage was ≅20x for H animals and 0x for L animals (S3 Fig). A recent study identified a small deletion, followed by a repetitive insertion, within the *nhj-1* gene (termed *nhj-1(vv148)*) that is present in a subset of ostensibly wild-type strains of *C. elegans* (S4A Fig) [45]. PCR-based analyses showed that *vv148* was present in L animals, but not H animals (S4B-E Figs). To test if *nhj-1(vv148)* was causative for regulating Tc1 mobility in the soma, we used CRISPR/Cas9 to generate an independent null allele in the *nhj-1* gene in H wild-type animals. The resultant allele (termed *gg875*) is predicted to be a strong loss-of-function for *nhj-1* as it introduces a premature stop codon and throws the majority of *nhj-1* out of frame (S4A Fig). *nhj-1(gg875)* animals exhibited radically reduced Tc1 mobilization rates in the soma, to a similar degree as *nhj-1(vv148)* (Fig 3B). The data establish that loss of NHJ-1 radically lowers the rate of Tc1 mobilization from *Tc1::mScarlet* in soma tissues.

### NHJ-1 acts in the soma to limit somatic Tc1 mobilization

To ask if NHJ-1 functions in the soma to regulate Tc1 mobilization, and not indirectly via the germline, we asked if animals lacking germ cells still exhibited altered Tc1 mobilization rates in the soma. Indeed, *glp-1(q224ts)* animals, which lack 99% of their germ cells when grown at the non-permissive temperature of ≥25℃ [46], exhibited rates of Tc1 mobilization similar to those of animals with intact germlines (Fig 3C). Additionally, loss of NHJ-1 did not obviously affect the rate at which Tc1 mobilized in germ cells (Table 1). We conclude that one function of NHJ-1 is to regulate Tc1 mobility, specifically in the soma.

### NHJ-1 is a general repressor of DNA transposons

We next asked if the loss of NHJ-1 might suppress mobilization of other transposons. We quantified Tc1 mobilization in *nhj-1(vv148)* animals harboring the *Tc1::sfgfp* reporter gene. The analysis showed far fewer sfGFP positive cells forming in the soma of *nhj-1(vv148)* animals than in *nhj-1(+)* animals (Fig 4A). We next quantified mobilization rates for two endogenous Tc1 elements in *nhj-1* mutant and *nhj-1(+)* animals using a quantitative PCR-based assay (S5 Fig). We observed that endogenous Tc1 elements were ≅1000 fold less likely to have mobilized in *nhj-1* mutants than in wild-type animals (Fig 4B). Similar effects were observed in *glp-1(q224ts)* animals lacking germ cells, indicating that NHJ-1 regulates endogenous Tc1 elements in the soma (Fig 4C). Finally, Tc3 is another Mariner class DNA transposon, which possesses little sequence homology to Tc1 [47]. Endogenous Tc3 mobilization rates were 10-fold lower in *nhj-1* mutant animals than in *nhj-1(+)* animals (Fig 4B). We conclude that loss of NHJ-1 is a general suppressor of transposon mobility in the *C. elegans* soma.

**Fig 4.**
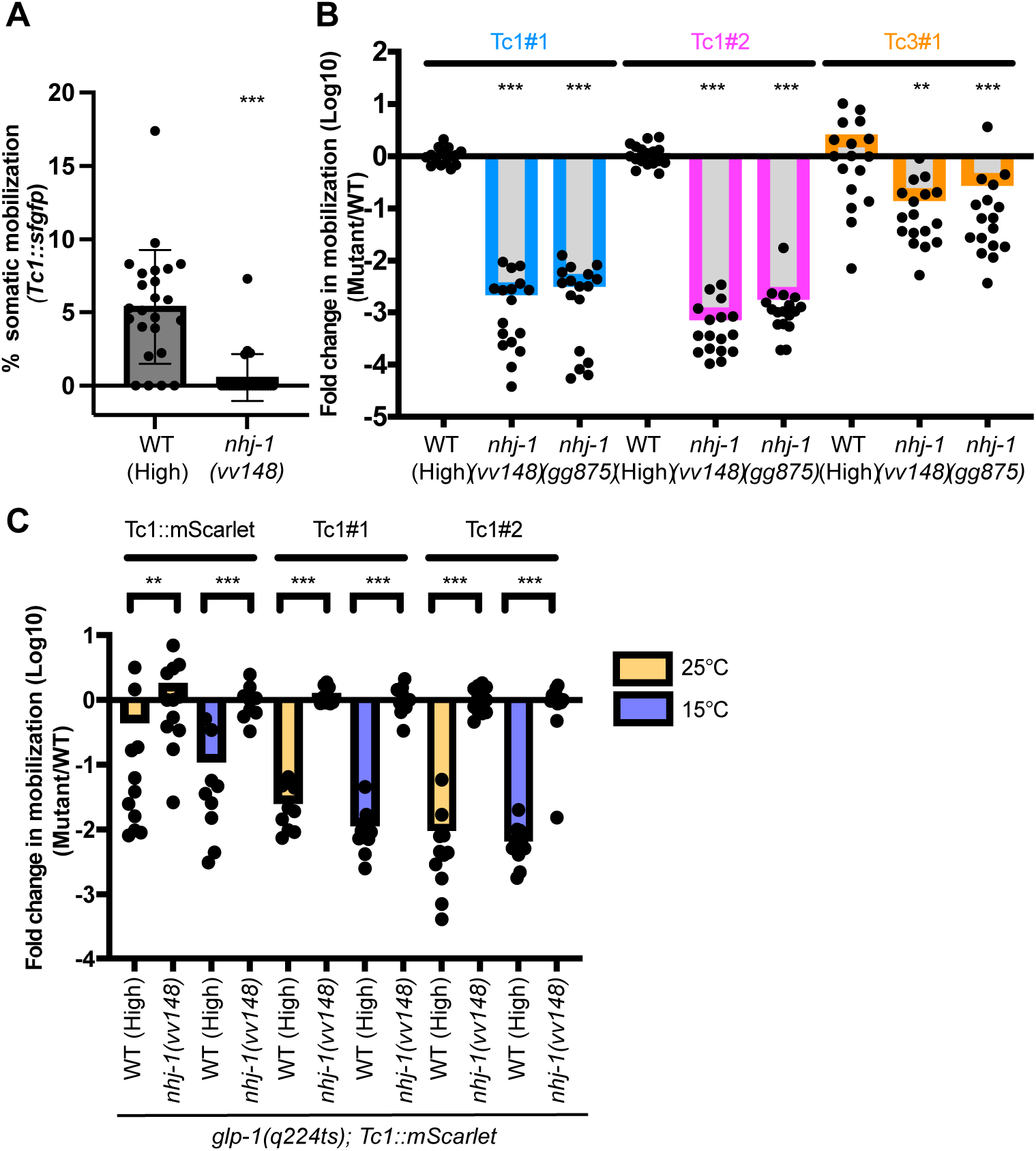
*nhj-1* is a general suppressor of transposon mobility. (A) *nhj-1* suppresses Tc1 mobilization from the *Tc1::sfgfp* reporter gene. Scoring for *Tc1::sfgfp* was conducted as described for *Tc1::mScarlet* in the Fig 2C legend, with the exception that scoring was done on an inverted microscope. Data-points are from at least three independent experiments scored on three different days. Error bars show standard deviations of the mean. ***p<0.001 (Mann-Whitney test). (B) qPCR-based quantification of transposon mobilization from two endogenous Tc1 insertion sites (Tc1 #1 Wormbase ID WBTransposon00000020 and Tc1 #2 WBTransposon00000033) and one endogenous Tc3 insertion site (Tc3 #1 WBTransposon00000062). **p<0.01; ***p<0.001 (Kruskal-Wallis test with Dunn’s test for multiple comparisons.) (C) qPCR based quantification of Tc1 mobilization from *Tc1::mScarlet* and Tc1# 1/2, using *glp-1(q224ts)* animals grown at 15°C or 25°C. **p<0.01; ***p<0.001 (Mann-Whitney test). (B-C) qPCR strategy used to quantify transposon mobilization in these panels is outlined in S5 Fig. Y axis; relative rates of transposon mobility expressed as the fold change in mutant lines relative to wild-type (High) in log10. Data-points are from at least three independent experiments.

### NHEJ increases transposon mobilization rates in somatic cells

How might NHJ-1 promote Tc1 mobilization? The non-homologous end joining (NHEJ) pathway repairs DNA breaks by directly ligating broken DNA ends. Three core components of the NHEJ machinery, LIG-4, CKU-80, CKU-70, are conserved in *C. elegans* [48, 49]. Importantly, previous studies find that *C. elegans nhj-1* acts in a genetic pathway with *lig-4*, *cku-80*, and *cku-70* to repair DNA double-strand breaks induced by chemotherapeutic agents and ionizing radiation, suggesting that NHJ-1 is a component of the *C. elegans* NHEJ DNA repair pathway [45,50,51]. Thus, the ability of NHJ-1 to promote transposon mobilization in the soma may relate to its function in NHEJ-based DNA repair. To test this idea, we asked if animals harboring loss-of-function mutations in *lig-4*, *cku-80*, or *cku-70* exhibited lowered rates of Tc1 mobilization in the soma, as described above for *nhj-1*. We used quantitative PCR (qPCR) to monitor Tc1 mobilization at two endogenous Tc1 elements and one Tc3 element in *lig-4*, *cku-80*, or *cku-70* mutants. *lig-4*, *cku-80*, and *cku-70* phenocopied *nhj-1* and caused Tc1 and Tc3 mobilization rates to decrease dramatically (Fig 5). The data show that during the normal course of growth and development of *C. elegans*, the use of NHEJ by somatic cells to repair their transposon-induced DNA breaks radically increases the rate of somatic transposon mobilization.

**Fig 5.**
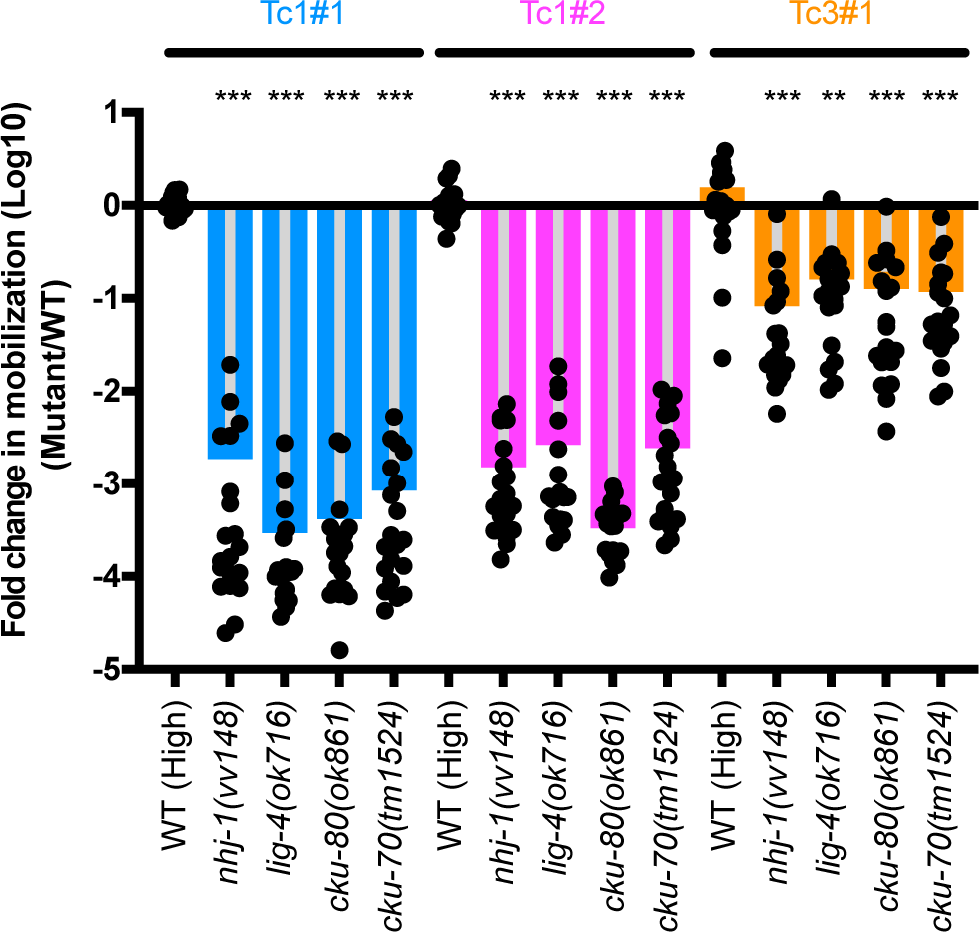
NHEJ repairs most somatic transposon mobilization events. The NHEJ factors LIG-4, CKU-80, and CKU-70 increase transposon mobility in the *C. elegans* soma. qPCR-based detection of Tc1 or Tc3 mobilization events from three endogenous transposable elements (see Fig 4 legend for details) in animals of the indicated genotypes. Y axis; relative rates of transposon mobility expressed as the fold change in mutant lines relative to wild-type (High) in log10. Data points are from at least three independent experiments. **p<0.01; ***p<0.001 (Kruskal-Wallis test with Dunn’s test for multiple comparisons).

## Discussion

Here we show that *C. elegans* lacking the NHEJ DNA repair machinery show a dramatic decrease in transposon mobilization (*i.e.* empty transposon sites) in somatic tissues. Thus, during normal growth and development, repair of transposon-induced DNA breaks by NHEJ in the soma is responsible for generating the vast majority of empty transposon sites accumulating in this tissue. By contrast, transposon-induced DNA breaks are repaired by HR DNA repair systems in the *C. elegans* germline [9]. Because HR uses homologous chromosomes (or sister chromatids) as templates for repair, HR will restore an excised transposon back into its original chromosomal position as a natural consequence of repair [9]. In other words, the vast majority of transposon excision events occurring in germ cells are masked by HR-based DNA repair. Because NHEJ repairs broken DNA by directly ligating DNA ends, NHEJ-based repair fixes (does not mask) transposon excision events. And because disabling NHEJ lowers the rate of somatic transposon mobilization rate to levels near that observed in the germline (Tables 1 and Fig 3), our data argue that the differential rates of transposon mobility observed between the *C. elegans* soma and germline can be explained largely, if not entirely, by the fact that these two tissue types use different DNA repair pathways to fix transposon-induced DNA breaks. Interestingly, homology-based repair and NHEJ-based repair are known to be preferentially used to repair *P* element transposon-induced DNA breaks in the germline and soma of flies, respectively. And NHEJ is known to repair *Sleeping Beauty* transposon-induced breaks in mammalian somatic cells [52–57]. Thus, tissue-specific differences in DNA repair pathway choice may influence transposon mobilization rates in other eukaryotes, in addition to *C. elegans*.

How and why might *C. elegans* use NHEJ in the soma, but not the germline, to help fix transposon-induced chromosome breaks? Previous studies document physical and/or genetic interactions between NHEJ factors and transposon-encoded proteins. For instance, the mammalian NHEJ factor Ku70 interacts physically with the transposase encoded by the *Sleeping Beauty* transposon [55]. Also, mammalian RAG proteins, which are widely considered domesticated transposases, interact with NHEJ factors (Ku70 and Ku80) to drive V(D)J recombination [58]. Finally, in *Paramecium*, NHEJ and the transposase-like PiggyMac cooperatively mediate programmed DNA elimination in developing macronuclei [59]. Thus, direct physical interactions between NHEJ factors and transposon-encoded proteins may explain how *C. elegans* deploys NHEJ to repair transposon-induced DNA breaks in the soma. Asking if Mariner transposases associate with NHEJ factors, and if this association only occurs in somatic cells, will be an important test of the model. A related question is why is NHEJ used to repair transposon damage in the soma but not the germline? HR can trigger large-scale chromosomal rearrangements when non-allelic templates are accidentally used for repair, and the risk of such non-allelic templating is likely to increase at or near repetitive elements, such as transposons. On the other hand, NHEJ does not depend on sequence homology for repair and, consequently, is unlikely to be negatively impacted by repetitiveness. Therefore, we propose that animals have evolved to use NHEJ to repair transposon-induced breaks in the soma in order to minimize chromosomal rearrangements that can lead to mitotic catastrophe and disease. And because homologs are paired in the germline, the risk of non-allelic templating is mitigated, allowing germ cells to use the highly accurate HR systems to fix their transposon-induced chromosomal damage.

While our data do not support the idea that transposons excise at very high rates uniquely in the soma as previously thought, our data do confirm that transposons are capable of excision in somatic cells of *C. elegans*, albeit at rates similar to those of the germline. Interestingly, we find that the systems limiting transposon mobilization in the germline (perinuclear foci and nuclear RNAi) do not limit mobilization in the soma. The data hint that other systems dedicated to taming transposons within the soma await discovery. The reporter genes described here should be useful in this regard.

## Materials and Methods

### Strains

WT N2, YY2002 ggSi29 [eft-3p::Tc1::(NLS)mScarlet(NLS)::tbb-2 3’UTR]; glp-1(q224ts), YY2003 ggSi29; glp-1(q224ts); nhj-1(vv148), YY2004 ggSi29; nhj-1(vv148), YY2044 rde-3(ne3370); ggSi41 [rpl-28p::Tc1::(NLS)sfgfp(NLS)::unc-54 3’UTR], YY2053 nhj-1(vv148), YY2054 ggSi55 [eft-3p::Tc1::(NLS)mScarlet(NLS)::tbb-2 3’UTR], YY2155 nhj-1(gg875), YY2096 rde-3(ne3370); ggSi55, YY2159 nhj-1(gg875); ggSi55, YY2185 mut-16(mg461); ggSi55, YY2186 mut-16(pk710); ggSi55, YY2177 znfx-1(ne4583); ggSi55, YY2187 deps-1(bn124); ggSi55, YY2191 ggSi41, YY2189 nrde-2(gg873); ggSi55, YY2192 ggSi41; nhj-1(vv148), FX1524 cku-70(tm1524), JK4605 glp-1(q224ts), RB873 lig-4(ok716), RB964 cku-80(ok861), VC40641 Million Mutation Project strain. All strains were cultivated on standard Nematode Growth Medium (NGM) plates and maintained at 20°C with Escherichia coli strain OP50 as food source unless otherwise noted [60]. JK4605 glp-1(q224ts) animals were maintained at 15°C. CRISPR strains were made with the co-CRISPR strategy [61] or Saptrap [62] combined with self-excising cassette selection [63]. Guide RNAs were chosen according to the CRISPOR.org tool [64]. Because the nhj-1(vv148) allele could be present in any strain, we made sure the mutant strains used for assaying Tc1 mobilization phenotypes (rde-3, mut-16, deps-1, znfx-1, nrde-2, cku-70, cku-80, lig-4) did not have the vv148 allele by PCR genotyping. gg873 allele, a putative null allele of nrde-2, was made by CRISPR and changes the 24th amino acid of nrde-2 from tyrosine to a stop codon [42].

### Plasmids

*Tc1::mScarlet* and *Tc1::sfgfp* reporter plasmids for CRISPR/Cas9 injection to make Tc1 reporter strains were constructed by using Saptrap to assemble components into the final pDD379 vector [62]. *Tc1::mScarlet* reporter plasmid donor sequence composes of Tc1 (1613 base pairs, including a TA dinucleotide at the 3’ end) inserted after the TA dinucleotide at position −7 to −8 of the 622 base pair *eft-3* promoter, followed by *mScarlet* sequence with nuclear-localization sequences (NLS) (SV40 NLS at 5’ end; *egl-13* NLS at 3’ end) and the *tbb-2* 3’UTR. *Tc1::sfgfp* reporter plasmid donor sequence composes of a 523 base pair *rpl-28* promoter, followed by a start codon and an 18 base pair flexible linker before the Tc1 sequence (1615 base pairs, including TA dinucleotide at both 5’ and 3’ ends). The Tc1 sequence is followed by another 18 base pair flexible linker before the *sfgfp* sequence (SV40 NLS at 5’ end; *egl-13* NLS at 3’ end of *sfgfp*) and the *unc-54* 3’UTR. The nuclear localized *sfgfp* sequence does not include a start codon. Tc1 sequences in both *Tc1::mScarlet* and *Tc1::sfgfp* donor plasmid constructs are the sequence of Tc1 with WormBase ID WBTransposon00000015, which does not contain any polymorphisms [5]. *Tc1::mScarlet* reporter plasmid guide RNA and homology arm components were designed to insert the reporter at Chr I: −5.32 cM. *Tc1::sfgfp* reporter plasmid guide RNA and homology arm components were designed to insert the reporter at Chr III: −0.85 cM. These sites were chosen according to insertion sites established for Mos1 mediated single copy insertion (MosSCI) that allow germline expression of inserts [65, 66].

### Microscopy

Larval stage animals were immobilized with 0.05% sodium azide and mounted on glass slides. Images were taken using the Axio Observer.Z1 fluorescent microscope (Zeiss) with the Plan-Apochromat 20x/0.8 M27 objective or the Plan-Apochromat 63x/1.4 Oil DIC M27 objective. Images were acquired using ORCA-Flash4.0 V2 digital camera (Hamamatsu) and ZEN 2 (blue edition) software (Zeiss). Images were processed using Fiji [67]. Microscope images in Fig 1B were spliced together from two images because a single image was not able to capture the full dimension of the animal.

### Soma Tc1 mobilization rate quantification

For the *Tc1::mScarlet* reporter, larval stage 4 animals were singled onto 6 cm NGM plates and progeny allowed to grow at 20°C. Four days later, each plate was examined under the AxioZoom.v16 fluorescence microscope (Zeiss) with the ApoZ 1.5x/0.37 objective. Animals were visually scored for presence or absence of somatic cells with mScarlet signal, to determine the percentage of progeny animals on a single plate that had somatic mScarlet signal. For the *Tc1::sfgfp* reporter, larval stage 4 animals were singled onto 6 cm NGM plates and progeny allowed to grow at 20°C. Three days later, animals were washed off the plate with M9 buffer, immobilized with 0.05% sodium azide, and mounted on glass slides. Slides were examined under the Axio Observer.Z1 fluorescent microscope (Zeiss) with the Plan-Apochromat 20x/0.8 M27 objective and animals were visually scored for presence or absence of somatic cells with sfGFP signal to determine the percentage of progeny animals on a single plate that had somatic sfGFP signal. For *glp-1* experiments, animals were bleach-synchronized and grown from embryonic stage at 25°C or 15°C until they reached larval stage 4. At larval stage 4, the plates were examined under the AxioZoom.v16 fluorescence microscope (Zeiss) with the ApoZ 1.5x/0.37 objective and animals were visually scored for presence or absence of somatic cells with mScarlet signal to determine the percentage of animals on each plate that had somatic mScarlet signal. All visual scoring of soma fluorescence was done with the examiner blinded to genotype. For the soma Tc1 mobilization rate quantified in Table 2, *Tc1::mScarlet* animals (YY2054) were bleach-synchronized and grown from embryonic stage at 20°C until larval stage 4. At larval stage 4, animals were observed for somatic Tc1 mobilization by mScarlet signal under the AxioZoom.v16 fluorescence microscope (Zeiss) with the ApoZ 1.5x/0.37 objective.

### Germline Tc1 mobilization rate quantification

Larval stage 4 animals were singled onto 6 cm NGM plates and allowed to propagate at 20°C. 20x concentrated OP50 was added to plates during the interval of growth. After several days, at either F1 or F2/3 generation, each plate was examined under the AxioZoom.v16 fluorescence microscope (Zeiss) with the ApoZ 1.5x/0.37 objective and scored for presence or absence of germline Tc1 mobilization events, presenting as animals in which all cells are mScarlet signal positive. The generation at which animals are scored for Tc1 mobilization (F1 or F2/3) varies with genotype and depends on how frequent germline Tc1 mobilization events are in that particular genotype. Number of animals on each plate was estimated by counting animals in one region of the plate, then multiplying by plate area. Germline Tc1 mobilization rates were calculated using the Poisson distribution formula [10, 28]: f = -(ln(N/T))/n, f: germline Tc1 mobilization frequency, T: total number of plates scored, N: total number of plates without germline Tc1 mobilization events, n: average number of animals per plate. n is calculated by summing the number of animals on each plate for a genotype, then dividing this sum by the total number of plates for that genotype. This calculation method addresses the concern of counting a single germline Tc1 mobilization event multiple times (jackpot effect) [10, 28]. In genotypes where no germline Tc1 mobilization event was detected, germline Tc1 mobilization rates were indicated as less than the rate that would have been if one plate of all plates scored had germline Tc1 mobilization event(s). All visual scoring was done with the examiner blinded to genotype.

### Whole genome sequencing and analysis

*C. elegans* genomic DNA was extracted using Gentra Puregene Tissue Kit (Qiagen, #158667) and DNA library preparation was performed using NEBNext Ultra II FS DNA Library Prep with Sample Purification Beads Kit (NEB, #E6177). Whole-genome sequencing was performed using an Illumina NextSeq instrument, with an Illumina NextSeq Mid-Output 300-cycle kit to obtain paired-end 150 bp reads at 30x coverage (Biopolymers Facility, Harvard Medical School). The quality of the library was assessed using *fastqc* (https://www.bioinformatics.babraham.ac.uk/projects/fastqc/). Paired-end reads for each library were mapped to the reference genome of *C. elegans* (PRJNA13758, WBcel235) using *bwa mem* v0.7.17 (http://bio-bwa.sourceforge.net/bwa.shtml). Compatible SAM files were merged, then sorted into bam files using *samtools* v1.3.1 [68]. The BAM file was indexed to create BAI files using *samtools* to load onto Integrative Genomics Viewer (IGV) [69]. The BAM and BAI files were loaded onto IGV and manual inspection identified a location with unmapped reads in the gene *nhj-1/scb-1* in L strains, but not in the H strain.

### Tc1 mobilization footprint analysis

*ggSi29*; *glp-1*(*q224ts*) strain (YY2002) animals were bleach-synchronized and grown from embryonic stage at 25°C for 3 days to reach adult stage. Adult worms were washed off the plate with M9 buffer and lysed with worm lysis buffer (50mM KCl, 10mM pH8.3 Tris, 2.5mM MgCl_2_) with proteinase K (0.2 mg/ml). Lysates were used as templates for PCR reactions using primers: *eft-3*pTc1mSc F: CTACCGTCCGCACTCTTC; *eft-3*pTc1mSc R: CTTACGCTTCTTCTTTGGC. PCR products were run on an agarose gel, and band size around 100 base pairs was excised and gel purified (Qiagen #27206). Purified PCR products were TA-cloned using the pGEM-T Easy Vector System (Promega, A1360), according to the manual instructions. White transformants were isolated, grown in liquid culture, and miniprepped. Plasmids were sent for Sanger sequencing using the primer M13F-40: 5’-GTT TTC CCA GTC ACG AC-3’.

### Tc1 mobilization qPCR assay

Larval stage 4 animals were singled onto 6 cm NGM plates and progeny allowed to grow at 20°C. Four days later, whole plate lysates were obtained by washing worms off with M9 buffer and lysing with worm lysis buffer (50mM KCl, 10mM pH8.3 Tris, 2.5mM MgCl_2_) with proteinase K (0.2 mg/ml). DNA lysates were used for qPCR assays using the iTaq Universal SYBR Green Supermix (Bio-Rad, 1725121), as instructed by the manual, and run on CFX Connect Real-Time PCR Detection System (Bio-Rad, 1855201). qPCR conditions were as follows: Initial denaturation at 95°C for 5 min followed by 40 cycles of denaturation at 95°C for 15 sec and annealing/extension/plate read at 61°C for 30 sec. Tc1 mobilization fold changes were calculated using the comparative Cq method [70]. Median Cq value of wild-type strain with H phenotype was used as the baseline for calculating Tc1 mobilization fold change for each experiment. *eft-3* was used as the housekeeping gene. If Cq value was undetectable by the end of the 40-cycle qPCR run, Cq value of 41 was used for the fold change calculation. For *glp-1* experiments, animals were bleach synchronized and grown from embryonic stage at 25°C or 15°C. When animals reached larval stage 4, whole plate lysates were obtained and used for qPCR as described above. Primers used for qPCR analysis are as follows: *eft-3* F: 5’-GTGAACGTGGTATCACCATC-3’; *eft-3* R: 5’-CGTACCAGTGATCATGTTC-3’; Tc1#1 F: 5’-CCGATCATCAATCATAGCG-3’; Tc1#1 R: 5’-GAGAACATTTGTGCGAG-3’; Tc1#2 F: 5’-GCGAGAAAAGGTATACTC-3’; Tc1#2 R: 5’-GTAGACATTATGCACCATTC-3’; Tc3#1 F: 5’-CGATTACAGAAGCCATCC-3’; Tc3#1 R: 5’-GGTCTCATCAGTAGTGACTC-3’. Tc1#1 refers to the Tc1 transposon with Wormbase ID WBTransposon00000020. Tc1#2 refers to the Tc1 transposon with Wormbase ID WBTransposon00000033. Tc3#1 refers to the Tc3 transposon with Wormbase ID WBTransposon00000062.

### Statistical analysis

Somatic Tc1 mobilization rate visual scoring and Tc1 mobilization qPCR assays were analyzed with Mann-Whitney test (for one comparison) or Kruskal-Wallis test with Dunn’s test for multiple comparisons (for more than one comparison) using GraphPad Prism version 9.2.0 (www.graphpad.com).

## Acknowledgements

We thank past and previous members of the Kennedy lab for helpful discussions and comments. We thank the BPF Genomics Core Facility at Harvard Medical School for their expertise and instrument availability that supported this work. Some strains were provided by the CGC, which is funded by NIH Office of Research Infrastructure Programs (P40 OD010440).

## Author Contributions

Conceptualization: C.C., S.K.; Methodology: C.C., S.K.; Investigation: C.C., D.J.P.; Formal analysis: C.C., D.D.L., D.J.P.; Writing: C.C., S.K.; Supervision: S.K.

---

**S1 Fig.**
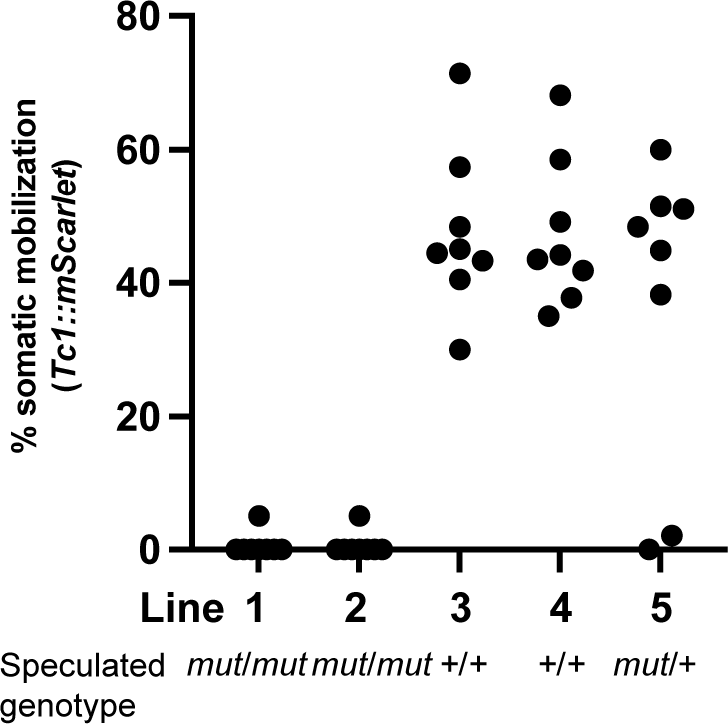
A genetic cross between ostensibly wild-type animals with Low and High somatic Tc1 mobilization phenotypes suggests the presence of a mutation that represses somatic Tc1 mobilization in H animals. Five progeny were isolated from a cross between Low and High animals and used to establish lineages (Lines 1-5). Eight animals from each line were singled to a plate and progeny were scored for the percentage of animals with bright mScarlet somatic cell signal. The results are consistent with a recessive mutation causing a low somatic Tc1 mobilization phenotype in L animals.

**S2 Fig.**
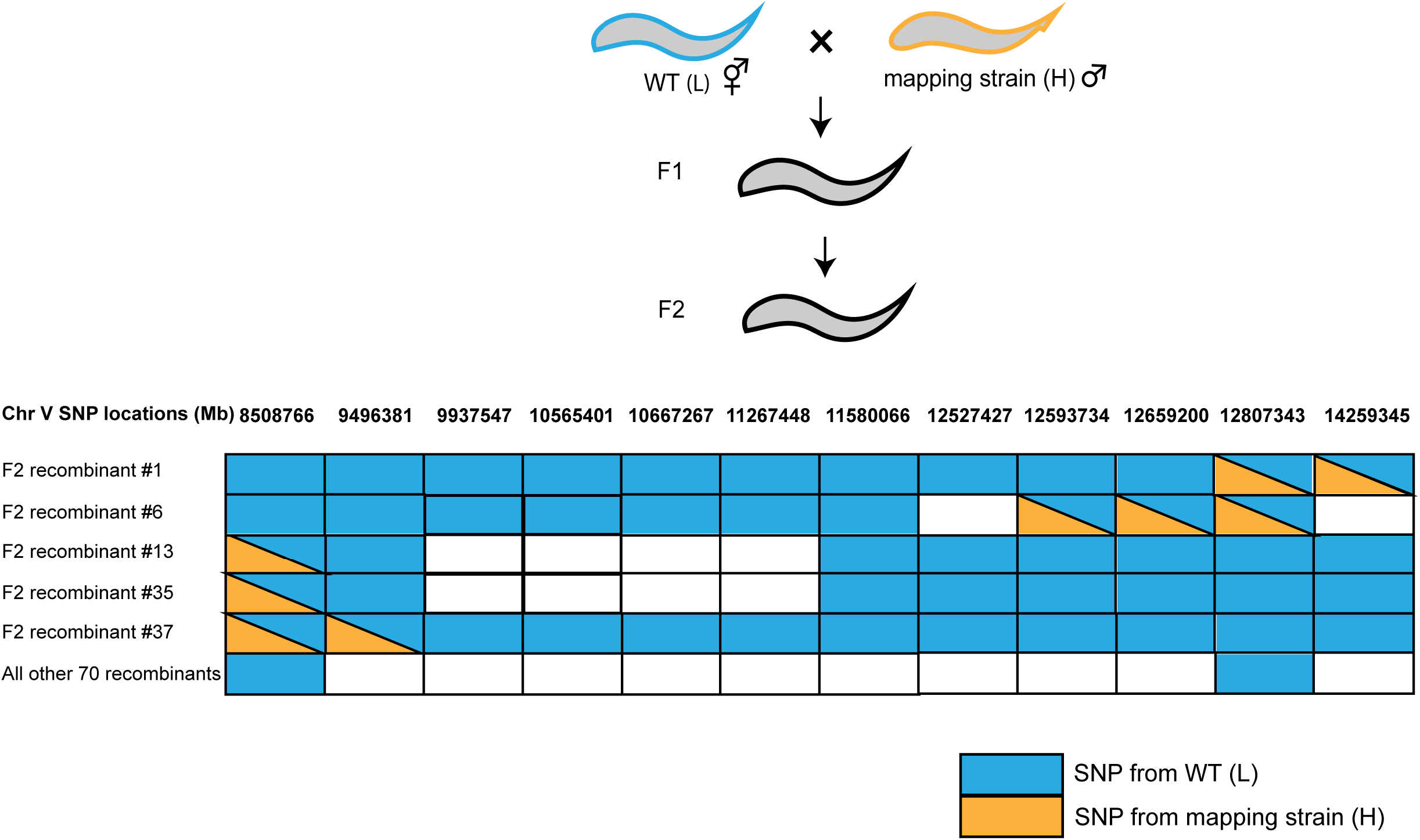
Positional mapping to identify location of allele causing low soma Tc1 mobilization phenotype. A genetic cross was performed between a Low somatic Tc1 mobilization strain and a strain from the Million Mutation Project (VC40641), which exhibits a High phenotype. 75 F2 progeny with the L phenotype were isolated and genotyped using SNPs present in VC40641. The results map the causal mutation to a region between 9.4 and 12.6 Mb on chromosome V. Five informative recombinants are shown. The remaining 70 F2 animals were homozygous for SNP markers of the L chromosome at and surrounding this region. Blue indicates SNP from the wild-type L chromosome and orange indicates SNP from VC40641(mapping strain) chromosome.

**S3 Fig.**
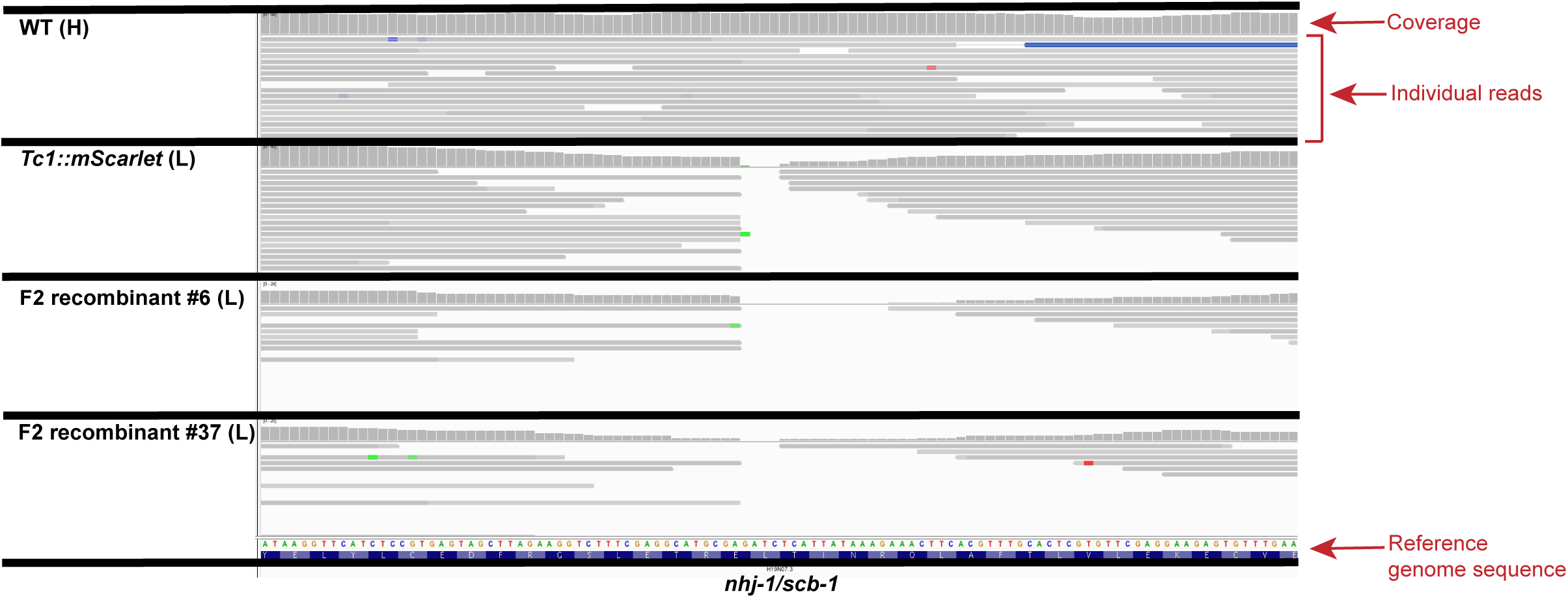
Lack of whole-genome sequencing coverage in L animals in a region of the *nhj-1* gene. Screenshots from Integrative Genomics Viewer (IGV) showing read coverage in H and L animals from whole genome sequencing data (horizontal gray bars) aligned to the reference genome. Note the absence of sequence coverage in L, but not H, sequencing data in the interval shown, which corresponds to the coding section of the *nhj-1* gene. F2 recombinants #6 and #37 are from the mapping cross described in Fig S2.

**S4 Fig.**
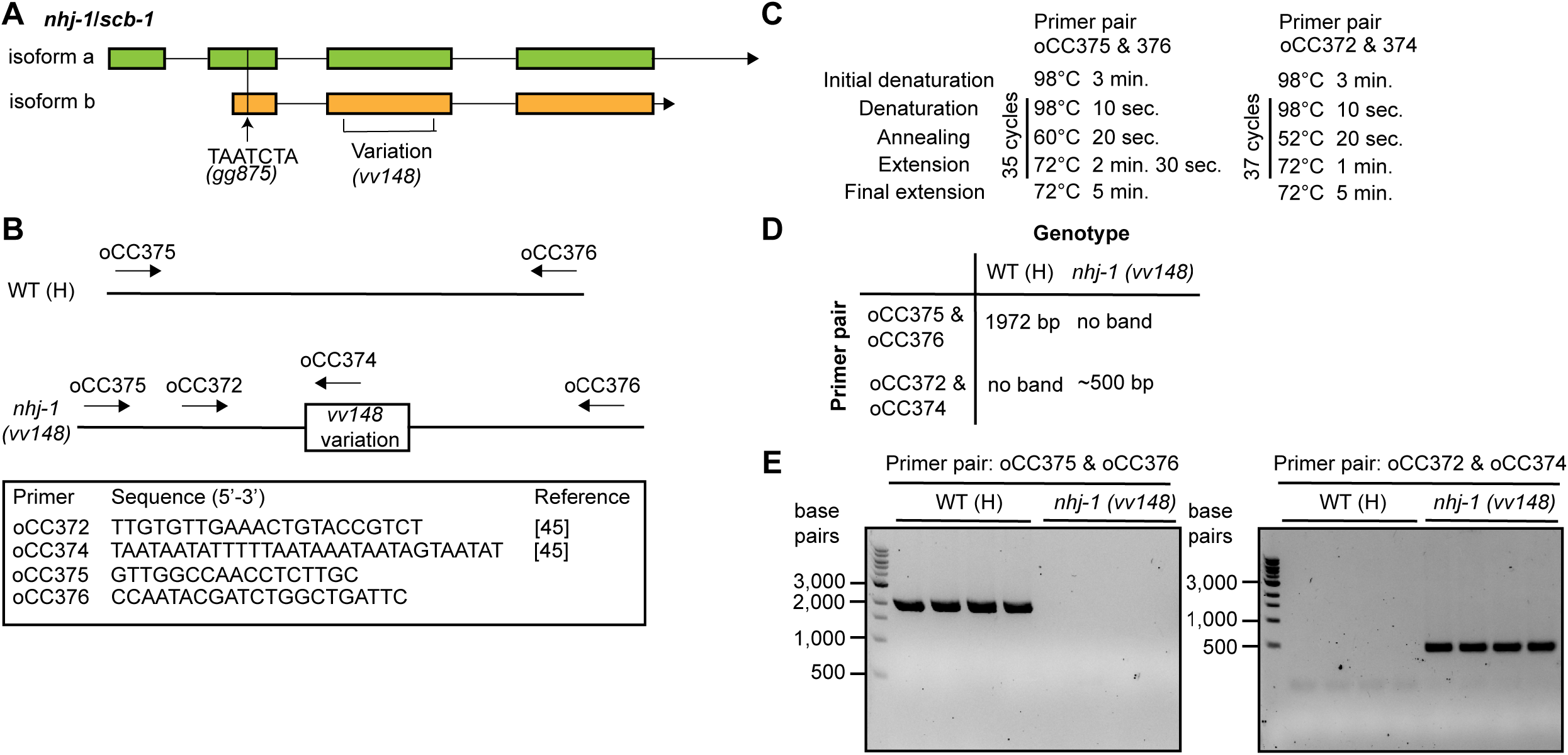
Description of *nhj-1* alleles from this study and genotyping information for *nhj-1(vv148)*. (A) *vv148* is a repetitive insertion/deletion allele (described in [45]). *gg875* is an insertion of 7 nucleotides (TAATCTA) near the 5’ end of *nhj-1*. (B) Genotyping primers for *vv148*. Primers oCC375 and oCC376 anneal on either side of the *vv148* variation, and will amplify only if *vv148* is not present. Primer oCC374 lies within the *vv148* variation [45], and will only amplify with oCC372 if *vv148* is present. (C) Genotyping conditions for *vv148*. (D) Expected outcomes for indicated PCR reactions for wild-type (H) and *nhj-1(vv148)* animals. (E) The indicated PCR products from wild-type (H) and *nhj-1(vv148)* animals run on an agarose gel and stained with ethidium bromide.

**S5 Fig.**
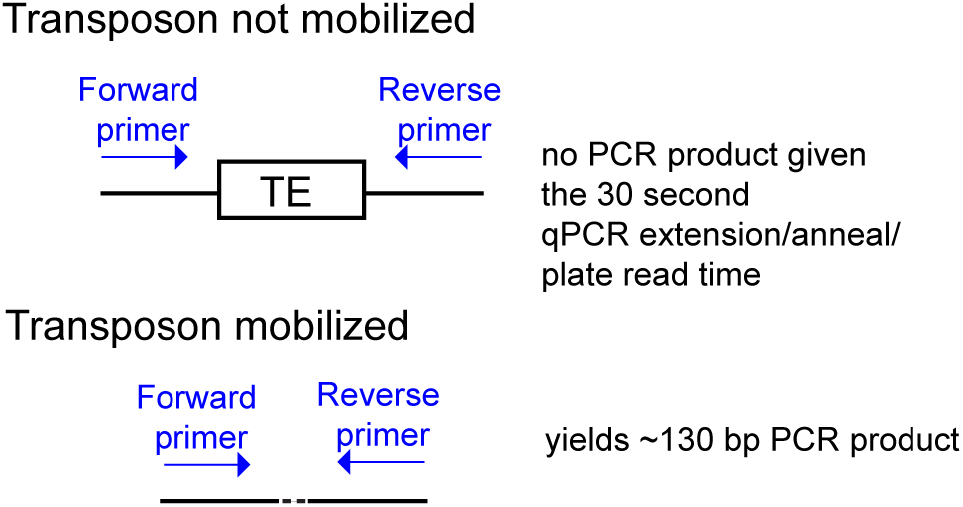
qPCR-based assay to quantify Tc1 and Tc3 mobilization. Primers are designed to flank transposons, and a short qPCR annealing/extension/plate read time (30 seconds) is used, so that only chromosomes where Tc1 has mobilized are efficient templates for PCR. TE, transposable element.

